# Galaxy External Display Applications: Closing a dataflow interoperability loop

**DOI:** 10.1101/642280

**Authors:** Daniel Blankenberg, John Chilton, Nate Coraor

## Abstract

Interoperability of datasets, tools, and resources is essential to modern scientific investigation and analysis. The necessity to gather disparate datasets together, perform analysis with a collection of discrete tools, and visualize the results remains a standard approach for exploring and making sense across scientific research domains. Here, we describe the Galaxy External Display Application (GEDA) framework which provides researchers with the ability to facilitate the interoperability of Galaxy user data and external resources, while promoting findability, accessibility, and reuse. The only requirement on the external resource for GEDA accessibility is that it is able to accept a parameter value that contains a URL pointing to user data.

## Introduction

Interoperability of datasets, tools, and resources is essential to modern scientific investigation and analysis. The necessity to gather disparate datasets together, perform analysis with a collection of discrete tools, and visualize the results remains a standard approach for exploring and making sense across scientific research domains^1,2^.

We have previously described the ability of Galaxy^3–6^ to ingest datasets from external data warehouse resources^7^, such as the UCSC Table Browser^8^, the EMBL-EBI European Nucleotide Archive (ENA; https://www.ebi.ac.uk/ena/about), various InterMine servers^9^, and others, through the use of DataSource tools. These DataSource tools enable researchers to easily mix-and-match data from any number of available resources directly into a Galaxy workspace. Once datasets have been loaded into the user’s workspace, they are able to configure and execute a wide-range of analysis tools. While, in many cases, the static outputs of bioinformatic tools are sufficient to generate tables, graphs, and meaningful results, there are often subsequent, and intermediate, next steps that often involve visualization.

Galaxy has built-in visualization capabilities^10^ that enable the building of various chart-types, circos-style genome-wide viewers, interactive phylogenetic trees, as well as custom genome browsers. Although enabling several visualization abilities, a significant drawback with these approaches is that these systems require the development, deployment, and use of code that is customized for use within the Galaxy platform. This can pose a formidable barrier to developers and can limit the reusability and accessibility of these facilities.

There exists a growing collection of standalone web-servers that provide visualization and analysis capabilities within discrete resources. Many of these web-servers and stand-alone applications, e.g. UCSC Genome Browser^11^, GBrowse^12^, IGV^13^, InterMine^9^, IGB^14^, IOBIO^15^, etc., allow users to upload their own datasets directly via submission from their computer, or by providing a URL to a web-hosted copy of the data. In cases when a dataset is local to a researcher’s computer, simply uploading the file directly can be the easiest approach, however, there can be significant drawbacks, including connectivity, transfer speeds, and the potential waste associated with downloading a dataset from one location to simply upload to another. Especially for large datasets, providing a URL link is often advantageous, however it does come with the difficulty associated with requiring a user to have access to, and knowledge of how to operate, web-hosting services.

## Results and Discussion

Here, we describe the Galaxy External Display Application (GEDA) framework which provides researchers with the ability to facilitate the interoperability of Galaxy user data and external resources, while promoting findability, accessibility, and reuse. The only requirement on the external resource for GEDA accessibility is that it is able to accept a parameter value that contains a URL pointing to user data. GEDAs that are available for a particular Galaxy dataset will appear as labeled links within the expanded preview of each particular dataset. Clicking on the link within the user’s dataset will open a new browser window and forward the user to the external resource, along with a customized URL pointing to the dataset contents or a dynamically generated manifest describing the dataset and its location. While this approach is often utilized to display and visualize user data, the external resource can be an analysis application or even data deposition service -- any service that accepts a URL parameter value is interoperable with Galaxy through the use of GEDAs.

GEDAs are declared using a straightforward XML-based description (figure 1) and associated with datatypes (figure 1.B), allowing or disallowing hierarchical inheritance across datatypes, as specified. The design of GEDAs are simple, yet highly extensible. A GEDA consists of a `“display”` tagset, that contains one or more `“link”` definitions, with each link having a `“url”` defined, along with a set of `“param”`eters that can be declared. In the simplest cases, they can be defined statically (see figure 1), with hard-coded resource URLs, that simply define a placeholder for a dataset URL. The GEDA framework will automatically generate a unique URL for the dataset to be passed to the external resource. GEDAs can also be dynamically generated (figure 2, 3, and 4), with links and options coming from externally managed flat files or through the Galaxy Data Table configuration system^16^. This enables a GEDA to be customized and updated with new options without requiring changes to the XML definition or the Galaxy codebase. Various filters (figure 3 and 4) can be applied to a GEDA to restrict access to Galaxy datasets that match defined criteria, such as belonging to a specific organism, genome build, and other metadata values. In cases where a GEDA is not available for a specific dataset and configuration, any otherwise potential links will not be created. By only displaying resources that are accessible for a particular dataset, resource findability is maximized.

**Figure 1:**
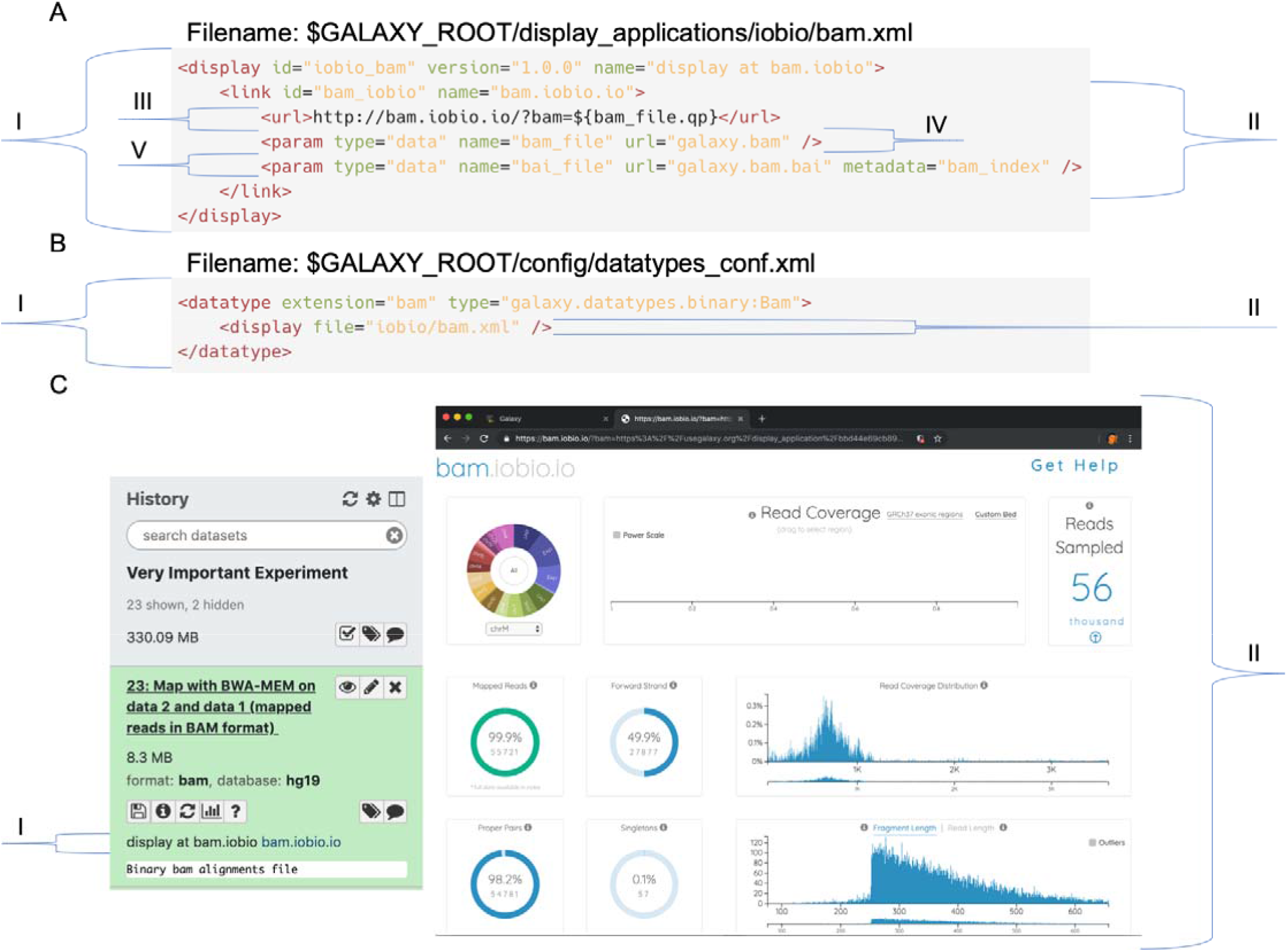
Anatomy of a basic Galaxy External Display Application. (A) A GEDA definition file contains several elements that describe how to provide interoperability with external resources, with (A.I) defining the entire application, by providing an `“id”`, `“version”`, and display `“name”`. Here, the display application description displayed to the user will start with “display at bam.iobio”. (A.II) Every GEDA has one or more `“link”` elements which contain a user actionable unit, as defined by an `“id”` and a `“name”` that will be used as the clickable display element for the resources. (A.III) A URL template is created that can utilize placeholders (e.g. bam_file) that will be populated by the GEDA framework and used as the URL to forward the user to. Here, a qp (quote_plus) method is being applied to the URL, that contains the BAM file data, to perform URL query string encoding. (A.IV) A parameter of type `“data”` is being created from the Galaxy dataset of interest, this parameter can be referenced within the URL, or in subsequent parameters (see figure 3). The value specified within the `“url”` attribute (here: “galaxy.bam”) will be used as the terminal “filename” being passed to the external resource. (A.V) BAM files within Galaxy have a file-based metadata element named “bam_index” that is also available for the external resource to fetch and use, which has been named as “bai_file” internally to the GEDA. Here, the resource-facing filename is defined as “galaxy.bam.bai” at the same base URL as the BAM file. (B) To instruct Galaxy to load the GEDA we defined in (A) for the BAM datatype (B.I), we add a `“display”` tagset (B.II) that references the GEDA XML file within the datatypes_conf.xml file. (C) The user-experience of accessing a GEDA is shown through the creation of a link (C.I) “display at <u>bam.iobio bam.iobio.io</u>”, that will send the user in their browser to the (C.II) external resource along with a URL to the Galaxy dataset content.

**Figure 2:**
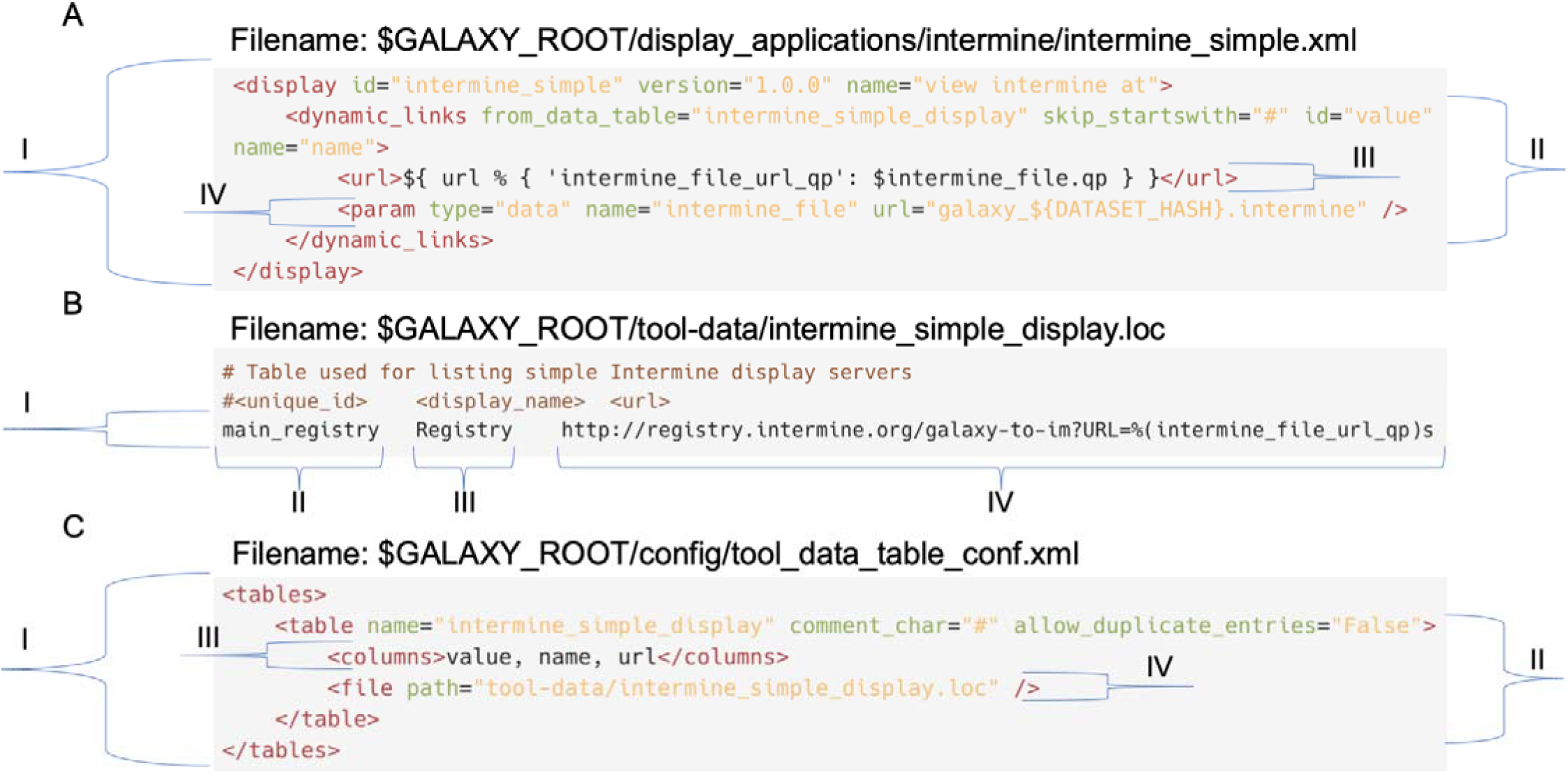
Anatomy of a dynamic Galaxy External Display Application. (A.I) A dynamic GEDA is initially defined in the same fashion as the static type (see figure 1). (A.II) A set of `“dynamic_links”` are defined according to the entries contained within a Galaxy Data Table (see panels B and C), with the link `“id”` and displayed `“name”` coming from the `“value”` and `“name”` fields, respectively. (A.III) This URL is generated using the Python string format operator (%) using the value of `“url”` from the Data Table, along with a dictionary containing an encoded URL to the “intermine_file” dataset content. (A.IV) The URL containing the InterMine dataset content is defined as containing a “DATASET_HASH” value that is unique to the Galaxy instance. This can be helpful to differentiate between multiple datasets within external resources that may only utilize or display the base filename of uploaded content; Galaxy dataset names can also be used (figures 3 and 4). (B) A flat file (colloquially referred to as a “.loc” or “location” file) is tab-delimited and can be modified outside of the Galaxy codebase to configure this display application. There is a single entry shown (B.I) which contains 3 columns: (B.II) a unique id for the entry, (B.III) the display name for the entry, and (B.IV) a variable named `“url”` that contains the URL template used in (A). (C) The Data Tables configuration (C.I) can contain any number of defined `“table”s` (C.II) which declare a `“name”` for the table, as well as to ignore lines in the file that begin with a comment character (#) and to not allow duplicate entries. The (C.III) columns for the three fields in the table are defined as `“value”`, `“name”`, and `“url”`. The (C.IV) source of the data table content is loaded from the specified flat file (B).

Different analysis pipelines often create datasets of dissimilar datatypes, despite many of these formats containing equivalent information. Because there can be differences in format between the dataset as it exists in the user’s workspace, and that which is accepted by the external resource, we have integrated GEDAs with Galaxy’s datatype conversion system (figure 3.VI). This allows datasets to be automatically converted on-demand to a derived dataset that is able to be consumed by the external resource. Additionally, any needed index or lookup table files can be created to enable fast, semi-random byte-range-based access to dataset content. For example, the GEDA for displaying VCF files at the UCSC Genome Browser server (figure 3) is defined to work for standard text-based VCF files, but when a user clicks the link to display the dataset at the server, job tasks are automatically configured and launched in the background to both compress the VCF with bgzip and to also build a Tabix index for the new compressed dataset. This generalized approach is able to facilitate accessibility, interoperability, and reusability of both the user data and the external (to Galaxy) resource.

**Figure 3:**
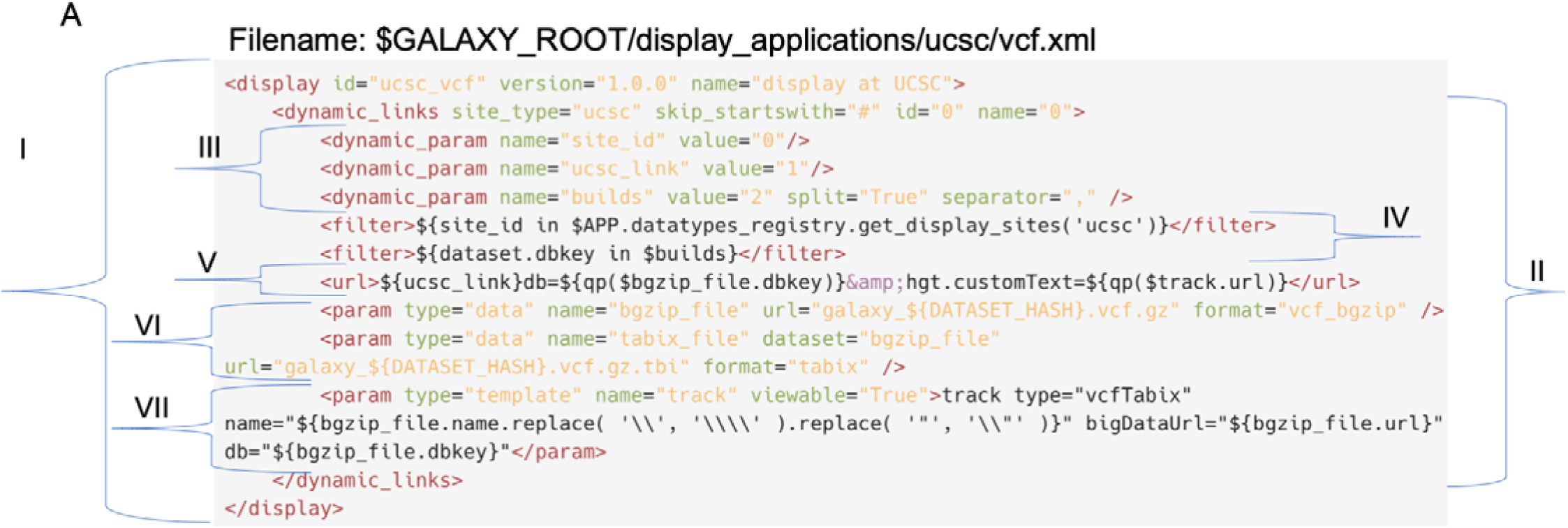
Anatomy of a dynamic Galaxy External Display Application utilizing filters and templates. (A.I) A dynamic GEDA is initially defined in the usual manner (see figure 2) for standard text-based VCF files. (A.II) A set of `“dynamic_links”` are defined according to the entries contained within an internal Galaxy system `“site_type”`. Additional (A.III) dynamic parameters are defined in a named fashion from the content of the site_type entry. The third parameter, “builds” is generated as a list from a comma-separated value within the site_type entry. (A.IV) Two filters are applied to determine if a GEDA link will be generated for the dataset. First to confirm that this UCSC Genome Browser site (e.g. main, mirror, or domain specific) has been enabled by the Galaxy Administrator. And, secondly, to confirm that this dataset belongs to a genome build that is valid for this particular UCSC Genome Browser site. Should any of the filters fail, then the GEDA will not be available and the link will not be generated for the user. The resource URL (A.V) is generated dynamically with the Galaxy content being defined using a UCSC Track definition file syntax ("track” template parameter) using the bigDataUrl mechanism. This GEDA is designed to work for standard VCF files, and (A.VI) we use Galaxy’s automatic datatype conversion system twice. First to bgzip the text-based VCF file and then to build a Tabix index on the bgzip VCF file. We (A.VII) define a template-type parameter named “track” that will be dynamically generated and presented as an externally viewable file to the external resource. This Track file provides the track type, a track name, dbkey (genome build), and a bigDataUrl value that references the bgzip VCF dataset content. This approach enables the UCSC Genome Browser to make efficient, semi-random access to the user’s dataset by only transferring the subset of data required for a particular user view.

GEDAs are not only limited to remotely hosted web-servers. For example, the popular Integrative Genomics Viewer (IGV) is available as a stand-alone Java-based desktop application that is executed locally on a user’s computer. The IGV software is able to load user datasets by direct file path loading and by pulling datasets from provided URLS. When the IGV GEDA (figure 4) is accessed via the local mode, the user is forwarded to a specific port (60151) bound by IGV on their computer via the localhost mechanism. The IGV desktop application will then load the datasets from the Galaxy server as needed, making proper use of available indexes to limit the amount of data that needs to be transferred at a time for any particular view. These complex, yet quintessential, details are all shielded from the user, of course, with the user experience consisting of clicking a link in Galaxy and having data loaded into their IGV desktop application.

**Figure 4:**
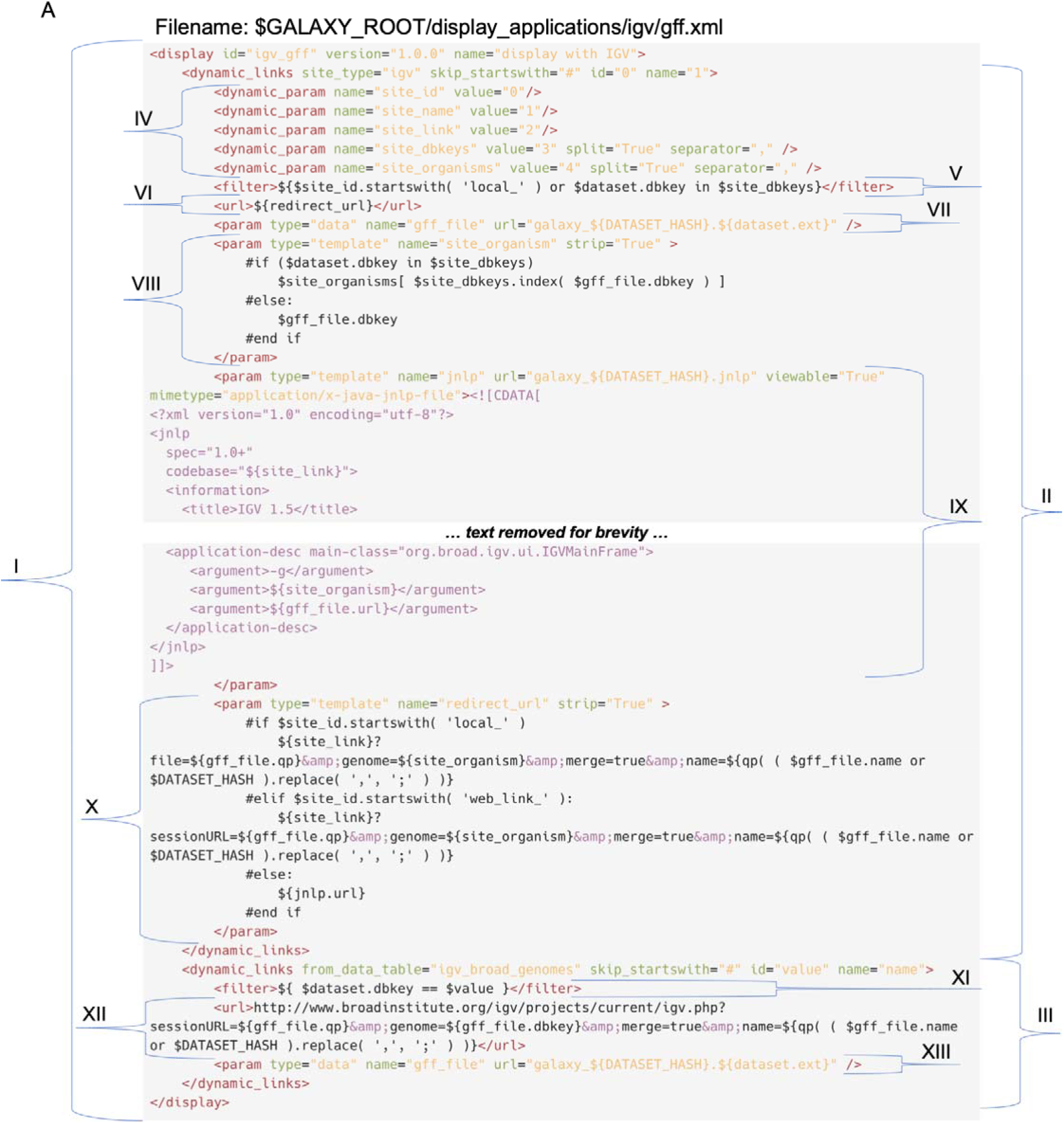
Anatomy of a complex Galaxy External Display Application interoperating with locally installed desktop software, which can be optionally launched over the web. This dynamic GEDA is initially defined in the usual manner (I), however this GEDA has two sets of `“dynamic_links”` tagsets (II and III), where each is able to generate multiple external resource links. The first link set (II) is generated from the Galaxy built-in sites list for IGV, with dynamic parameters (IV) defined similarly as in (3.III). GEDA links are filtered (V) to include sites designed for directly operating on running local software or where the genome build for the dataset is enabled at the external resource. The URL (VI) is dynamically generated from a template-based parameter named “redirect_url” (X). This GEDA will work for GFF datasets (VII) and derivatives, with the URL file extension coming directly from the specific hierarchical datatype for the dataset. A (VIII) template-based parameter named “site_organism” is used to determine the genome build ID valid for the IGV software from a look-up table or as directly specified, with the `“strip”` attribute indicating to remove any whitespace surrounding the populated parameter value. A (IX) template-based parameter named “jnlp” dynamically generates a Java Network Launch Protocol (JNLP) description file that can be used launch a new instance of the IGV desktop application with the user’s GFF dataset. The (X) redirect_url used for the external resource URL (VI) is created in one of three different versions, depending upon the type of external resource site definition (local, web, or jnlp). The second set of `“dynamic_links”` (III) is defined from a Galaxy Data Table that is populated from an external URL (http://igv.broadinstitute.org/genomes/genomes.txt), enabling automatic syncing with the latest IGV genome build support. A filter (XI) is applied to ensure that only datasets with genome builds available within this IGV database will have GEDA links created. A URL (XII) is created that forwards the user and dataset content URL directly to the launch IGV page at the Broad Institute, along with a defined track name and genome build. The GFF dataset content (XIII) is defined as in (VII).

## Conclusions

There are many computational resources available that allow users to upload their own data for use in analysis and visualization tools by providing URLs. These resources vary from genome browsers, to analysis pipelines and dataset dashboards, to locally running desktop applications. To facilitate streamlined findability, accessibility, interoperability, and reuse of these resources with the Galaxy platform, we have developed the Galaxy External Display Application framework. The GEDA framework enables effortless integration of Galaxy datasets and these disparate external computational resources. GEDAs can be defined statically, or abstractly, with context-specific dynamic interactivity. Regardless of the complexity of the GEDA, or the computations and actions occurring behind the scenes, the user experience remains simple, accessible, powerful, and consistent: user clicks link, user goes to the external resource along with a URL describing their dataset contents, remote resource loads user provided data. Currently, over 30 individual GEDAs have been developed, including configurations for Ensembl, GBrowse, IGV, IGB, InterMine, IOBIO, Rviewer, and the UCSC Genome Browser, with many having been contributed by the extended Galaxy community.

## Acknowledgements

The authors are grateful and indebted to the Galaxy team and the Galaxy community for all of their contributions.

## References

1. Blankenberg, D., Taylor, J. & Nekrutenko, A. Online resources for genomic analysis using high-throughput sequencing. Cold Spring Harb. Protoc. 2015, 324–335 (2015).

2. Qu, K. et al. Integrative genomic analysis by interoperation of bioinformatics tools in GenomeSpace. Nat. Methods 13, 245–247 (2016).

3. Afgan, E., Baker, D., Batut, B. & Van Den Beek, M. The Galaxy platform for accessible, reproducible and collaborative biomedical analyses: 2018 update. Nucleic acids (2018).

4. Giardine, B. et al. Galaxy: a platform for interactive large-scale genome analysis. Genome Res. 15, 1451–1455 (2005).

5. Blankenberg, D., Von Kuster, G. & Coraor, N. Galaxy: a web based genome analysis tool for experimentalists. Current protocols in (2010).

6. Blankenberg, D. & Hillman-Jackson, J. Analysis of next-generation sequencing data using Galaxy. Methods Mol. Biol. 1150, 21–43 (2014).

7. Blankenberg, D. et al. Integrating diverse databases into an unified analysis framework: a Galaxy approach. Database 2011, bar011 (2011).

8. Karolchik, D. et al. The UCSC Table Browser data retrieval tool. Nucleic Acids Res. 32, D493–6 (2004).

9. Kalderimis, A. et al. InterMine: extensive web services for modern biology. Nucleic Acids Res. 42, W468–72 (2014).

10. Goecks, J. et al. Web-based visual analysis for high-throughput genomics. BMC Genomics 14, 397 (2013).

11. Kent, W. J. et al. The human genome browser at UCSC. Genome Res. 12, 996–1006 (2002).

12. Stein, L. D. Using GBrowse 2.0 to visualize and share next-generation sequence data. Brief. Bioinform. 14, 162–171 (2013).

13. Thorvaldsdóttir, H., Robinson, J. T. & Mesirov, J. P. Integrative Genomics Viewer (IGV): high-performance genomics data visualization and exploration. Brief. Bioinform. 14, 178–192 (2013).

14. Freese, N. H., Norris, D. C. & Loraine, A. E. Integrated genome browser: visual analytics platform for genomics. Bioinformatics 32, 2089–2095 (2016).

15. Miller, C. A., Qiao, Y., DiSera, T., D’Astous, B. & Marth, G. T. bam.iobio: a web-based, realtime, sequence alignment file inspector. Nat. Methods 11, 1189 (2014).

16. Blankenberg, D., Johnson, J. E., Galaxy Team, Taylor, J. & Nekrutenko, A. Wrangling Galaxy’s reference data. Bioinformatics 30, 1917–1919 (2014).

